# An ascending vagal sensory-central noradrenergic pathway modulates retrieval of passive avoidance memory

**DOI:** 10.1101/2024.04.09.588717

**Authors:** Caitlyn M. Edwards, Inge Estefania Guerrero, Danielle Thompson, Tyla Dolezel, Linda Rinaman

## Abstract

**Background:** Visceral feedback from the body is often subconscious, but plays an important role in guiding motivated behaviors. Vagal sensory neurons relay “gut feelings” to noradrenergic (NA) neurons in the caudal nucleus of the solitary tract (cNTS), which in turn project to the anterior ventrolateral bed nucleus of the stria terminalis (vlBNST) and other hypothalamic-limbic forebrain regions. Prior work supports a role for these circuits in modulating memory consolidation and extinction, but a potential role in retrieval of conditioned avoidance remains untested.

**Results:** To examine this, adult male rats underwent passive avoidance conditioning. We then lesioned gut-sensing vagal afferents by injecting cholecystokinin-conjugated saporin toxin (CSAP) into the vagal nodose ganglia (Experiment 1), or lesioned NA inputs to the vlBNST by injecting saporin toxin conjugated to an antibody against dopamine-beta hydroxylase (DSAP) into the vlBNST (Experiment 2). When avoidance behavior was later assessed, rats with vagal CSAP lesions or NA DSAP lesions displayed significantly increased conditioned passive avoidance.

**Conclusions:** These new findings support the view that a gut vagal afferent-to-cNTS^NA^-to-vlBNST circuit plays a role in modulating the expression/retrieval of learned passive avoidance. Overall, our data suggest a dynamic modulatory role of vagal sensory feedback to the limbic forebrain in integrating interoceptive signals with contextual cues that elicit conditioned avoidance behavior.

## Background

Interoceptive information about the physiological state of the body is communicated continuously to the brain to shape current and future motivated behaviors (1). For example, visceral sensory signals arising in the gut can influence innate avoidance of stimuli that are naturally aversive (i.e., not requiring prior experience) (2). In addition, our lab recently reported that visceral sensory feedback can influence avoidance of conditioned stimuli that become aversive after being paired with an innately aversive experience (3,4). Sensory feedback about the body’s internal state is conveyed to the brain through blood-borne and neural pathways, the latter of which include vagal sensory pathways (5–9). In one study, rats with selective destruction of vagal sensory afferents displayed less avoidance of innately aversive stimuli, but displayed prolonged freezing responses to an auditory conditioned fear cue (10), suggesting that vagal sensory signaling can increase innate avoidance responses while reducing or suppressing learned fear responses. This discrepancy provides an interesting distinction between neural circuits that control innate avoidance/anxiety-like behavior and circuits that use experience and memory to generate conditioned fear responses. However, since sensory vagotomies in that study were performed prior to fear conditioning, it is unclear what aspect of fear learning was affected (i.e., acquisition, consolidation, retrieval, and/or extinction). To our knowledge, the necessity of vagal afferents and downstream circuits in the expression/retrieval of conditioned avoidance behavior has not been explored. While some research has examined the influence of vagus nerve stimulation on memory consolidation (11–13) and extinction (14–16), the potential role of vagal afferent signaling in the retrieval of passive avoidance memory is untested.

The central axon terminals of vagal viscerosensory afferents innervate postsynaptic neurons in the caudal nucleus of the solitary tract (cNTS), including neurons comprising the A2 noradrenergic (NA) cell group (17,18). cNTS^A2^ neurons project to a number of hypothalamic and limbic forebrain regions in which NA signaling modulates arousal and stress responses (19). For example, cNTS^A2^ neurons densely innervate the anterior ventrolateral bed nucleus of the stria terminalis (vlBNST) (20,21), which receives additional NA input from the caudal ventrolateral medulla (A1 cell group) (22,23). A2/A1 neurons that innervate the vlBNST are recruited by a variety of acute innate stressors (2,19,24–29), and lesioning A2/A1 inputs to the vlBNST attenuates both innate stress-induced avoidance (30) and hypophagia (31). The vlBNST itself appears necessary for the expression of conditioned freezing responses to contextual fear stimuli (32,33), and has been implicated in other forms of conditioned behavioral suppression (34,35).

We recently reported that cNTS^A2^ neurons and neurons within the NA terminal-rich region of the anterior vlBNST are activated to express the immediate-early gene product, cFos, in rats exposed to a conditioned context that previously was paired with an aversive footshock (3). The recruitment of these cNTS^A2^ and vlBNST neurons by a learned stressor implicates this ascending circuit in modulating conditioned avoidance behavior.

The present study was designed to test the hypothesis that a vagal sensory-to-cNTS^NA^-to-vlBNST circuit modulates the expression of learned passive avoidance behavior. In *Experiment 1*, we sought to selectively lesion gastrointestinal vagal afferents by injecting cholecystokinin (CCK)-conjugated saporin toxin (CSAP) bilaterally into the nodose ganglia (36,37). In *Experiment 2*, we sought to selectively lesion NA inputs to the anterior vlBNST by injecting saporin toxin conjugated to an antibody against dopamine beta hydroxylase (DbH) (DSAP) bilaterally into the vlBNST (30,31,38). In order to isolate lesion effects on the retrieval/expression of passive avoidance memory without impacting memory acquisition or consolidation, CSAP or DSAP lesions were made in rats *after* passive avoidance training.

## Methods

All experiments were conducted in accordance with the National Institutes of Health *Guide for the Care and Use of Laboratory Animals* and were reviewed and approved by the Florida State University Animal Care and Use Committee.

### Passive Avoidance Training and Testing

Adult male Sprague Dawley rats (Envigo; 250-300g body weight) were pair-housed in standard tub cages in a temperature- and light-controlled housing environment (lights on from 0400 hr to 1600 hr). Rats were acclimated to handling for three days, with free access to water and rat chow (Purina 5001). Rats underwent passive avoidance training in a novel light/dark shuttle box (Colbourn Instruments, Allentown), with training performed 4-6 hr after lights on. The light/dark box comprised a light (illuminated) chamber with clear plastic walls and a smooth white plastic floor, and a dark (non-illuminated) chamber with black plastic walls and a metal grid floor. Each chamber measured 25 x 25 cm, with 28 cm-high walls and a ceiling. The two chambers were separated by a metal divider wall with a guillotine door that could be opened and closed remotely. For passive avoidance training, rats were initially placed individually into the light chamber of the box and the dividing guillotine door was lifted immediately to allow access to the dark chamber. As expected, due to their innate aversion to the light and preference for the dark, rats very quickly entered the dark chamber (average latency = 19.47s). Upon entry into the dark chamber, the guillotine door was closed. After a 5s delay, rats received a single mild electric footshock (0.6mA; 1s). Rats remained in the enclosed dark chamber for 30s following footshock, and then were returned to their homecage. Cagemates were similarly trained on the same day.

At least two days following training, rats underwent surgery to lesion gut-sensing vagal afferents or brainstem NA neurons that innervate the vlBNST, as described below (*CSAP/DSAP Injections*). Three weeks after training (at least two weeks after surgery), rats were tested for passive avoidance retention. For this purpose, rats were placed individually into the light chamber of the light/dark box, the guillotine door was opened, and the latency of rats to fully enter the dark chamber was recorded, with a pre-set maximum latency of 900s (15 min). During the retention test, the guillotine door remained open and no footshock was administered. Rats were allowed to freely explore both the light and dark chambers during the 900s test, and total time spent within each chamber was recorded. After testing was complete, rats were returned to their home cages. Four rats in the DSAP experiment (n=2/group) were excluded from analysis due to the incorrect GraphicState protocol being used during testing.

### Nodose CSAP Injections

For the CSAP experiment, we used a highly specific vagal afferent lesioning agent to target gastrointestinal vagal afferents that express the gene encoding the CCK A receptor *(Cckar)* (36,39). The lesioning agent (CSAP) is a conjugate of sulfonated CCK and the ribosome-inactivating protein, saporin [CCK-SAP; Advanced Targeting Systems, Inc.]. Rats were anesthetized with intraperitoneal injection of a cocktail containing ketamine (100 mg/kg) and xylazine (10 mg/kg) and placed into a supine position. The left and right nodose ganglia was identified and isolated as previously described (36). CSAP was diluted to 250 ng/μl in 0.15M NaCl vehicle containing 0.1% Fast Green. CSAP (N= 11 rats) or Fast Green vehicle alone (N= 5 rats) was delivered (1µl bilaterally) into each nodose ganglion by pressure injection using a pulled glass capillary attached to a micromanipulator. Rats were allowed to recover from surgery and were tested for passive avoidance retention at least two weeks after surgery to allow time for effective lesioning (36).

### BNST DSAP Injections

As in our previous studies (30,31), we used a highly specific approach to lesion NA neurons projecting to the anterior vlBNST. The lesioning agent (DSAP) is a conjugate of a mouse monocloncal antibody to dopamine beta-hydroxylase (DbH) and the ribosome-inactivating protein, saporin [Anti-DbH-SAP; Advanced Targeting Systems, Inc.]. Rats (N = 25) were anesthetized by inhalation of isoflurane (1-3% in oxygen) and placed into a stereotaxic apparatus. The anterior vlBNST (located 0.35mm posterior, 2.3mm lateral, and 7.4mm ventral to bregma) was targeted using a 10-degree angle from vertical to avoid passing through the lateral ventricle and dorsal BNST. DSAP was diluted to 40ng/100nl in 0.15M NaCl vehicle containing 0.05% CTB, which does not interfere with DSAP lesion efficacy (30). DSAP (N= 11 rats) or CTB vehicle alone (N= 10 rats) was delivered (200nl bilaterally) into the avlBNST by pressure injection (20nl/min for 10 min) using a Hamilton syringe driven by a motorized injector (Stoelting Co.). The syringe was left in place for 10 minutes after each injection. Rats were allowed to recover from surgery and were tested for passive avoidance retention at least two weeks later, which is sufficient for effective lesioning (30,31,40–42).

### Histology

Rats were deeply anesthetized with pentobarbital sodium (39 mg/ml i.p., Fatal Plus Solution; Butler Schein) and transcardially perfused with saline (100mL) followed by 4% paraformaldehyde (500mL). Fixed brains were removed from the skull, post-fixed overnight at 4°C, then cryoprotected in 20% sucrose. Brains were blocked and sectioned coronally (35 μm) using a Leica freezing-stage sliding microtome. Tissue sections were collected in six serial sets and stored at -20°C in cryopreservant solution (43) until immunohistochemical processing.

Primary and secondary antisera were diluted in 0.1M phosphate buffer containing 0.3% Triton X-100 and 1% normal donkey serum.

For rats used in the nodose CSAP lesion experiment, heads were post-fixed overnight at 4°C after brain removal. Fixed nodose ganglia were then dissected out, extracted and cryoprotected in 20% sucrose. Nodose ganglia were sectioned at 20 μm using a Leica cryostat and transferred directly to charged microscope slides for RNAscope-based analysis of lesion extent.

### Verification and Characterization of Nodose CSAP Lesion

Quantification of left and right nodose ganglia lesion extent included analysis of 2-3 sections through each nodose ganglia from each rat in the CSAP experiment. Sections were processed to detect surviving (i.e., non-lesioned) neurons expressing mRNA for the CCK-A receptor (*Cckar*). For this, the Rn-CCK-AR probe (412091, Accession No. NM_012688.3) and RNAscope Fluorescent Multiplex Assay (323100, ACDBio, San Franscico, CA) were used. Sections were incubated in channel 1-specific HRP (323104) and Cy5-TSAP (1:1000, tyramine signal amplification plus, NEL744E001KT; PerkinElmer Waltham, MA, USA). After RNAscope labeling, slides were subsequently incubated in mouse monoclonal antiserum against the neuronal marker HuC/D (1:2500; ThermoFisher Scientific; A-21271; AB_221448) followed by Cy3-conjugated donkey anti-mouse IgG (1:500; Jackson ImmunoResearch). Sections were imaged at 20x using a Keyence microscope (BZ-X710). The number of HuC/D-positive neurons present within each section and the number containing labeled CCK-AR transcripts were counted, with counts normalized to derive cell density within each analyzed area. Count densities within the left and right nodose ganglia were then averaged across 2-3 analyzed sections in each rat, and then combined and averaged within each lesion group. Cell density counts from three nodose ganglia could not be quantified due to ineffective extraction (CSAP: n = 2; Veh: n = 1). For comparison, left and right nodose ganglia from two non-manipulated control rats were also processed and analyzed to reveal potential physical damage arising from CSAP or Veh injections.

### Verification and Characterization of DSAP Lesion

One set of tissue sections from each rat in the BNST DSAP experiment (with each set containing a complete rostrocaudal series of sections spaced by 210 μm) was incubated in a mouse monoclonal antiserum against DbH (1:50,000; Millipore, MAB308; AB_2245740) followed by biotinylated donkey anti-mouse IgG (1:500; Jackson ImmunoResearch). Sections were then treated with Elite Vectastain ABC reagents (Vector) and reacted with diamino-benzidine (DAB).

Sections through the cNTS containing the A2 cell group and VLM containing the A1 cell group (∼15.2-14.15 mm caudal to bregma) were imaged at 20x using a Keyence microscope (BZ-X710). The total number of DbH+ neurons were counted bilaterally in each region and averaged per section. Sections through the pontine locus coeruleus (LC; ∼10-10.2 mm caudal to bregma) were imaged at 10x. Due to extremely dense DbH labeling in the LC, it was not possible to visualize and accurately count individual DbH+ cells. Instead, a region of interest (ROI) encompassing the DbH labeling in LC was drawn using Image J bilaterally and the area of each ROI was measured and averaged across four LC ROIs (equivalent to bilateral LC in 2 tissue sections per rat).

### Statistics

Data were analyzed using GraphPad Prism. Measures of passive avoidance behavior during the retention test (i.e., latency to enter and time in the dark chamber) were analyzed using unpaired Student’s t-tests. Data quantifying *Cckar*-expressing nodose ganglia neurons for the CSAP experiment and DbH immunolabeling (cell counts in cNTS and VLM, area measurements in LC) for the DSAP experiment were analyzed using unpaired Student’s t-test. In addition, anatomical data were correlated with passive avoidance behavior within subjects using linear regression. An alpha level of 0.05 (p ≤ 0.05) was used as the criterion for considering group differences to be statistically significant. Estimation statistics were used to report effect sizes for both passive avoidance behavior and verification of CSAP and DSAP lesions (44,45).

## Results

### Experiment 1: CSAP Lesions Passive Avoidance Task

Unpaired t-tests indicated that compared to vehicle-injected controls, CSAP-lesioned rats displayed more avoidance of the shock-paired dark chamber as measured by a longer latency to enter [t(14) = 2.396; p = 0.0311] (Fig. 1A) and less total time spent within the dark chamber [t(14) = 2.381; p = 0.0320] (Fig. 1B). Examining the unpaired mean difference between CSAP- and vehicle-injected rats using estimation statistics indicated that CSAP rats took an average of 174s (2.9min) longer to enter the shock-paired dark chamber compared to vehicle controls [95%CI 5.64s, 290s], and spent an average of 83.5s (1.4min) less in the shock-paired dark chamber during the 900s retention test [95%CI 19.2s, 159s].

**Figure 1.**
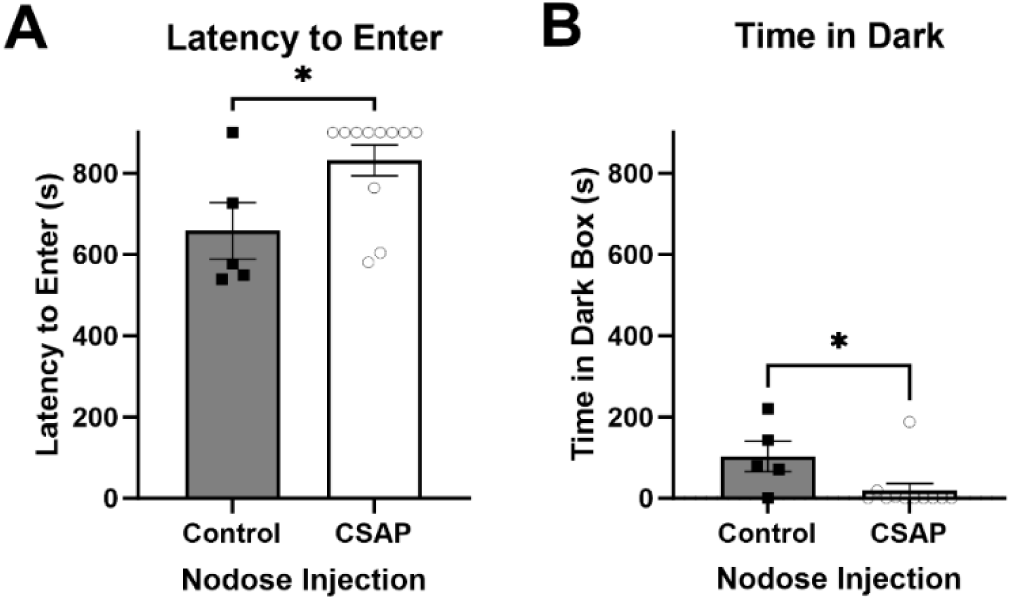
CSAP lesion increased passive avoidance behavior. (A) CSAP-lesioned rats took an average of 174s longer to enter the shock-paired dark chamber compared to vehicle-injected controls. (B) CSAP-lesioned rats spent an average of 84s less in the shock-paired dark chamber compared to vehicle-injected controls. *p < 0.05. Graphs display group mean +/- SEM, with individual data points plotted.

### Verification and Characterization of CSAP Lesion

One sample t-test indicated that the density of *Cckar*+ neurons in the left nodose of vehicle-injected and non-manipulated rats did not significantly differ (t(4) = 1.215; p = 0.2912).

Conversely, the density of left nodose *Cckar*+ cells was significantly reduced compared to the non-manipulated controls (t(8) = 14.63; p < 0.0001). Similar results were seen in the right nodose ganglia, in which the density of *Cckar*+ neurons in vehicle-injected rats was similar to that in non-manipulated rats (t(5) = 1.026; p = 0.3521), whereas the density of *Cckar*+ neurons in CSAP-injected rats was significantly lower than in non-manipulated controls (t(8) = 5.894; p = 0.0004).

When data from rats with nodose vehicle injections were compared to data from CSAP-injected rats, unpaired t-test confirmed effective lesion of *Cckar*-expressing neurons in the nodose ganglia of CSAP-injected rats [t(15) = 3.973; p = 0.0012] (Fig. 2G). Compared to vehicle-injected controls, the density of *Cckar*+ neurons was significantly reduced in the left nodose [t(12) = 4.890; p = 0.0004] (Fig. 2A), with a trending reduction in the right nodose [t(13) = 1.967; p =0.0709] (Fig. 2D). Examining the unpaired mean difference between CSAP- and vehicle-injected rats using estimation statistics indicated that CSAP rats had an average of 40 fewer *Cckar*+ cells/mm^2^ in the left nodose ganglion [95%CI 22, 57] and 30 fewer *Cckar*+ cells/mm^2^ in the right nodose ganglion compared to vehicle-injected controls [95%CI -3, 62].

**Figure 2.**
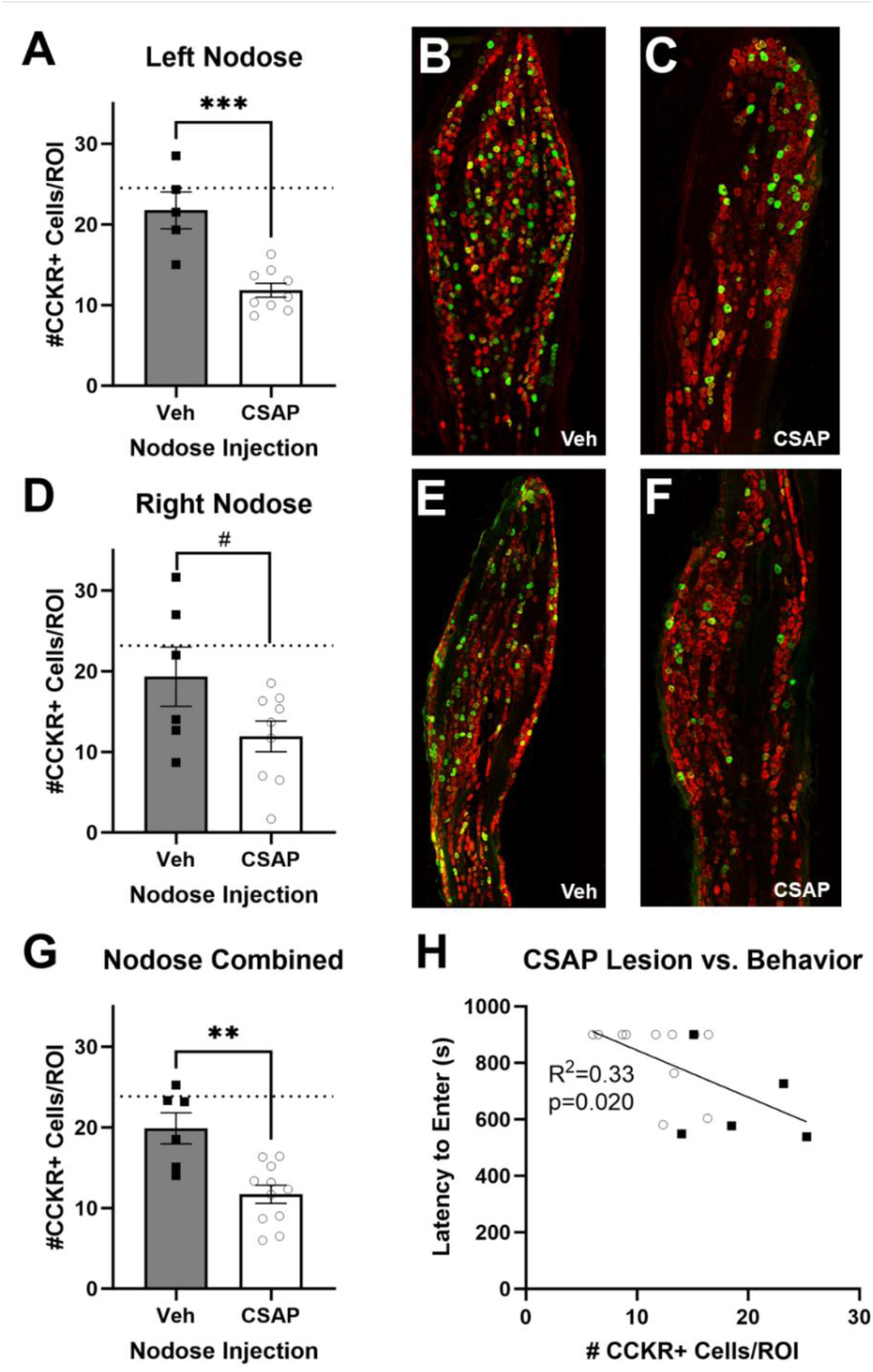
CSAP injection was effective at lesioning *Cckar*+ nodose ganglia neurons. (A) CSAP-lesioned rats displayed a 54.5% reduction in *Cckar*+ cells in the left nodose ganglia. (B+C) Representative images of *Cckar*+ (green) labeling and HuC/D (red) labeling in the left nodose ganglion of a vehicle control (B) and CSAP-lesioned rat (C). (D) Compared to vehicle-injected controls, CSAP-lesioned rats displayed 61.7% reduction in *Cckar*+ cells in the right nodose ganglia. (E+F) Representative images of *Cckar*+ (green) labeling and HuC/D (red) labeling in the right nodose ganglion of a vehicle control (E) and CSAP-lesioned rat (F). (G) Compared to vehicle-injected controls, CSAP-lesioned rats displayed 58.8% reduction in *Cckar*+ cells in the nodose ganglia combined. (H) Degree of CSAP lesion is negatively related to the latency to enter the dark chamber. #p < 0.10; **p < 0.01; ***p < 0.001. Graphs display group mean +/- SEM, with individual data points plotted. Dotted lines on graphs represents the average of two non-injected nodose ganglia.

We observed significant negative within-subjects relationships between the latency to enter the dark chamber and the extent of the CSAP lesion, including a negative correlation between the latency to enter and the number of *Cckar*+ cells in the combined left/right nodose ganglia [r(16) = -0.5760, p = 0.0195] (Fig. 2H) and in the left nodose ganglion alone [r(13) = - 0.5545, p = 0.0492] (Table 1). There was no significant correlation between latency to enter the dark chamber and the number of *Cckar*+ cells in the right nodose ganglion.

**Table 1:**
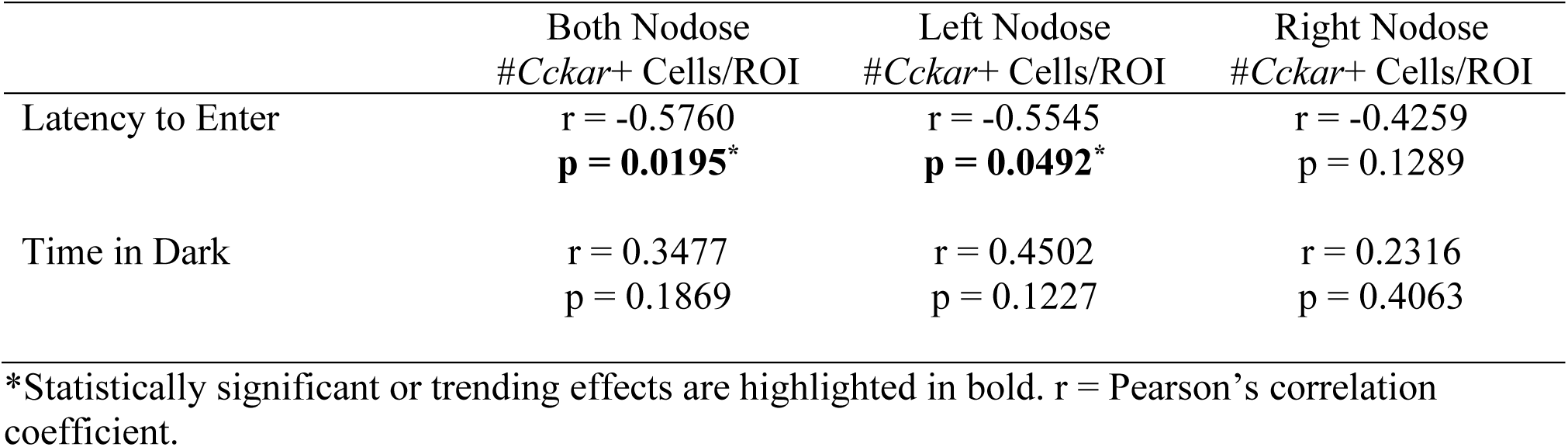
Correlations between the CSAP lesion and behaviors in the passive avoidance task.

|  | Both Nodose<br>#*Cckar*<sup>+</sup> Cells/ROI | Left Nodose<br>#*Cckar*<sup>+</sup> Cells/ROI | Right Nodose<br>#*Cckar*<sup>+</sup> Cells/ROI |
| --- | --- | --- | --- |
| Latency to Enter | $r = -0.5760$<br>**$p = 0.0195^*$** | $r = -0.5545$<br>**$p = 0.0492^*$** | $r = -0.4259$<br>$p = 0.1289$ |
| Time in Dark | $r = 0.3477$<br>$p = 0.1869$ | $r = 0.4502$<br>$p = 0.1227$ | $r = 0.2316$<br>$p = 0.4063$ |

\*Statistically significant or trending effects are highlighted in bold. $r$ = Pearson's correlation coefficient.

### Experiment 2: DSAP Lesions

#### Passive Avoidance Task

Unpaired t-test indicated that DSAP-lesioned rats displayed more avoidance of the shock-paired dark chamber as measured by a longer latency to enter [t(19) = 1.997; p = 0.0302] (Fig. 3A) and less time spent in the dark chamber [t(19) = 2.183; p = 0.0418] (Fig. 3B) compared to vehicle controls. Examining the unpaired mean difference between DSAP- and vehicle-injected rats using estimation statistics indicated DSAP rats took an average of 194s (3.23min) longer to enter the shock-paired dark chamber compared to vehicle controls [95%CI 4.08s, 380s], and spent an average of 160s (2.67min) less time within the shock-paired dark chamber during the 900s retention test [95%CI 33.7s, 322s].

**Figure 3.**
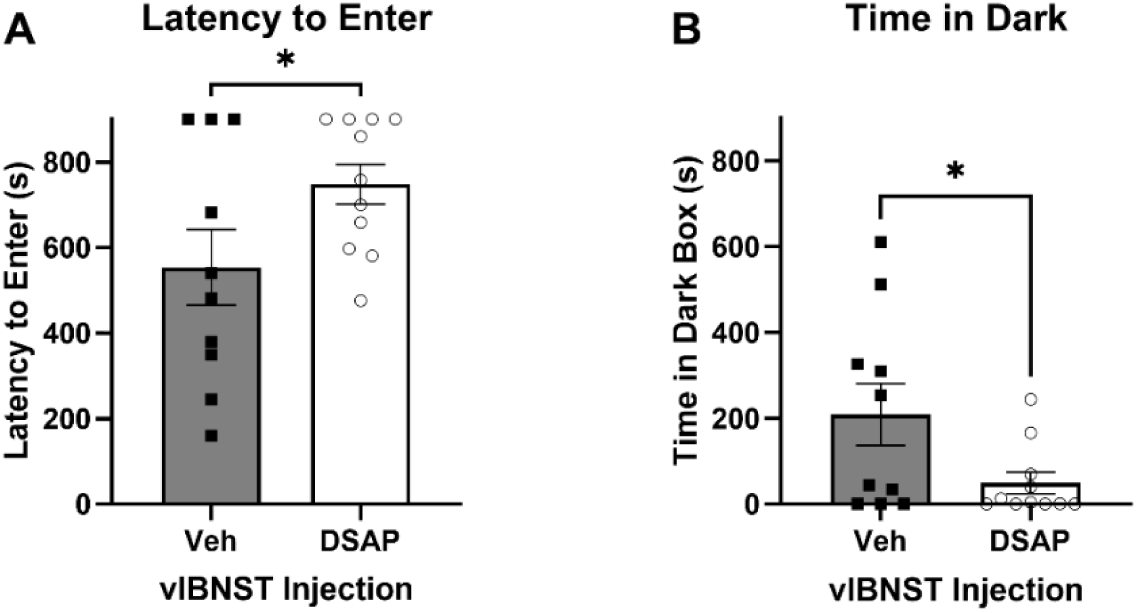
DSAP lesion increased passive avoidance behavior. (A) DSAP-lesioned rats took an average of 194s longer to enter the shock-paired dark chamber than vehicle-injected controls. (B) DSAP-lesioned rats spent an average of 160s less in the shock-paired dark chamber than vehicle-injected controls. *p < 0.05. Graphs display group mean +/- SEM, with individual data points plotted.

#### Verification and Characterization of DSAP Lesion

Unpaired t-test confirmed effective lesion of hindbrain DbH+ neurons. The number of DbH+ neurons per section was significantly reduced in both the cNTS [t(23) = 15.63; p < 0.0001] (Fig. 4A) and the VLM [t(23) = 10.30; p < 0.0001] (Fig. 4D). Examining the unpaired mean difference between DSAP- and vehicle-injected rats using estimation statistics indicated that DSAP rats had an average of 22.2 fewer DbH+ cNTS neurons per section [95%CI 19.5, 24.9] and 16.2 fewer DbH+ VLM neurons per section [95%CI 13, 18.9] compared to vehicle-injected controls, corresponding to an average loss of 60.1% and 47.1% of DbH+ neurons, respectively. Student’s t-test also revealed a moderate reduction in the area of DbH+ labeling within the LC in DSAP rats [t(23) = 3.00; p = 0.0062] (Fig. 4G). Examining the unpaired mean difference between DSAP- and vehicle-injected rats using estimation statistics indicated that the average ROI area in DSAP rats (i.e., DbH+ LC area) was 1.91mm^2^ smaller than in vehicle-injected controls [95%CI 0.715, 3.09], a reduction of approximately 19.3%. We observed similar depletion of DbH+ labeling within the LC in previous studies in which DSAP was injected into the vlBNST (30), likely due to sparse axonal inputs from the LC to the vlBNST and closely adjacent areas.

**Figure 4.**
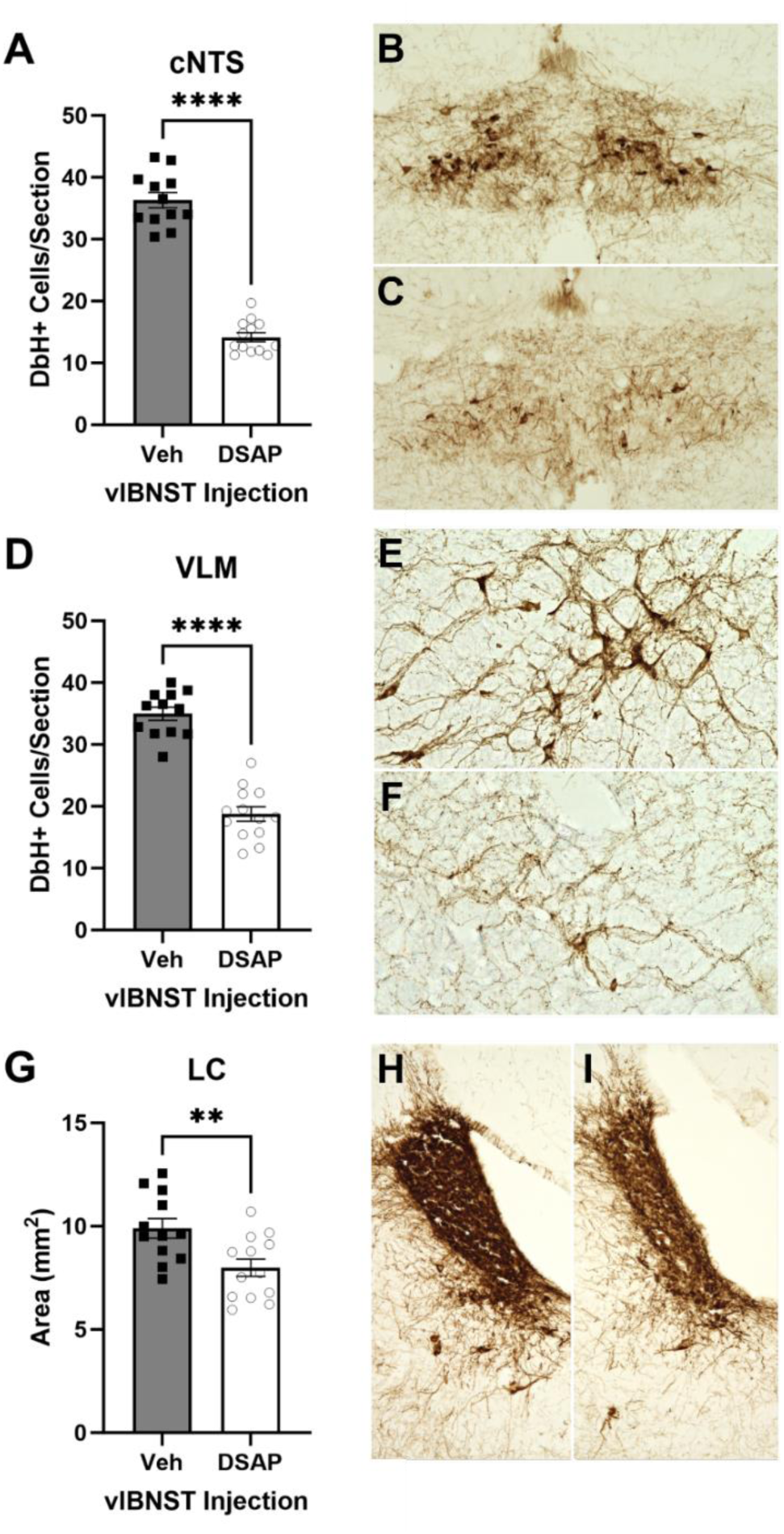
DSAP injection was effective at lesioning NA inputs. (A) Compared to vehicle-injected controls, DSAP-lesioned rats displayed a 60.1% reduction in DbH+ cells/section in the cNTS. (B+C) Representative images of DbH+ labeling in the cNTS of a vehicle control (B) and DSAP-lesioned rat (C). (D) Compared to vehicle-injected controls, DSAP-lesioned rats displayed 47.1% reduction in DBH+ cells/section in the VLM. (E+F) Representative images of DbH+ labeling in the VLM of a vehicle control (E) and DSAP-lesioned rat (F). (G) Compared to vehicle-injected controls, DSAP-lesioned rats displayed 19.3% reduction in DBH+ staining area in the LC. (H+I) Representative images of DbH+ labeling in the LC of a vehicle control (H) and DSAP-lesioned rat (I). **p < 0.01; ****p < 0.0001. Graphs display group mean +/- SEM, with individual data points plotted.

We observed negative within-subjects relationships between the latency to enter the dark chamber and the extent of the DSAP lesion, including a trending negative correlation between the latency to enter and the number of DbH+ neurons/section in the cNTS [r(19) = -0.4201, p = 0.0580] (Table 2) and a significant negative correlation between the latency to enter and the number of DbH+ neurons/section in the VLM [r(19) = -0.4467, p = 0.0423] (Table 2).

**Table 2:**
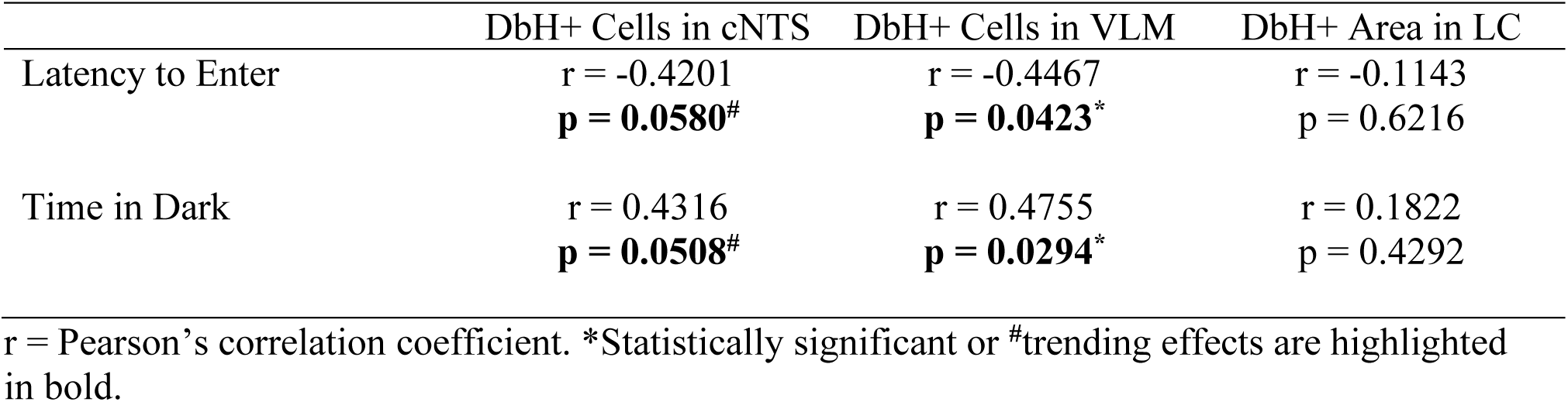
Correlations between the DbH immunolabeling and behaviors in the passive avoidance task.

|  | DbH+ Cells in cNTS | DbH+ Cells in VLM | DbH+ Area in LC |
| --- | --- | --- | --- |
| Latency to Enter | $r = -0.4201$<br>**$p = 0.0580^{\#}$** | $r = -0.4467$<br>**$p = 0.0423^*$** | $r = -0.1143$<br>$p = 0.6216$ |
| Time in Dark | $r = 0.4316$<br>**$p = 0.0508^{\#}$** | $r = 0.4755$<br>**$p = 0.0294^*$** | $r = 0.1822$<br>$p = 0.4292$ |

$r$ = Pearson's correlation coefficient. \*Statistically significant or $^{\#}$ trending effects are highlighted in bold.

Conversely, there was no significant correlation between latency to enter the dark chamber and the area of DbH+ staining within the LC.

## Discussion

Results from previous studies have implicated vagal afferent signaling and central noradrenergic signaling in avoidance behaviors in rodents (10,46–48). The present study is the first to demonstrate that expression of learned passive avoidance is modulated by vagal afferents and by noradrenergic neurons that provide axonal input to the vlBNST. Neurotoxin-induced loss of gut-sensing afferents in the vagus nerve or loss of noradrenergic inputs to the anterior vlBNST enhanced conditioned passive avoidance behavior in rats, as measured by their increased latency to enter and less time spent in the shock-paired dark chamber (Fig. 1, 3). The magnitude of these behavioral effects in *Experiments 1 and 2* was closely related to the extent of loss of *Cckar*+ nodose neurons in CSAP-treated rats (Table 1) and of DbH+ neurons within the cNTS and VLM in DSAP-treated rats, but was not related to the more modest loss of DbH immunolabeling within the LC in DSAP-treated rats (Table 2).

Most research into the influence of the vagus nerve on memory has focused on manipulations during memory consolidation and extinction (11–16). Electrical and pharmacological stimulation of the vagus nerve post-training (i.e., during memory consolidation) enhances passive avoidance memory in rodents (11,12). Interestingly, left cervical vagal nerve stimulation (VNS) in human patients during the consolidation phase immediately following learning enhances subsequent recognition memory (49). Consistent with our results indicating enhanced conditioned avoidance after vagal sensory lesions, VNS applied during retrieval *disrupts* memory performance in humans (50). Taken together, these data support the view that vagal afferent signaling plays a role in enhancing memory consolidation, while also contributing to disruption of memory retrieval.

The vagus nerve has been suggested to have indirect effects on memory by enhancing central noradrenergic signaling (51). Previous studies have shown that brain-wide depletion of norepinephrine enhances spatial, contextual, and avoidance memory retrieval in rats (52–54). Vagal afferents synapse directly onto A2 noradrenergic neurons in the cNTS (17,18), which provide the major source of noradrenergic input to the vlBNST (24,55). In *Experiment 2*, we demonstrate that lesioning noradrenergic inputs to the vlBNST enhances memory retrieval, providing circuit-specific insight regarding central sites at which norepinephrine depletion may influence memory retrieval. Thus, a vagal afferent-to-cNTS^NA^-to-vlBNST signaling pathway may serve to constrain or reduce fear memory retrieval, such that retrieval is enhanced when endogenous signaling through this pathway is interrupted by CSAP or DSAP lesions. It is important to note, however, that DSAP-induced destruction of NA neurons that innervate the vlBNST is accompanied by loss of other axon collaterals arising from the same NA neurons, which include axons in the hypothalamus (56). Loss of these additional NA pathways could contribute to the observed increase in conditioned passive avoidance behavior in DSAP rats.

An alternative interpretation of our experimental results is that the vagal afferent-to-cNTS^NA^-to-vlBNST pathway may be necessary to coordinate *active* (vs. passive) behavioral responses to threat, such that interruption of this circuit may bias the animal away from active and towards passive responses to threat. Consistent with this idea, while VNS reduces freezing behavior, a passive response to threat, VNS also has been shown to increase shuttling behavior in the active avoidance task (57). Furthermore, brain-wide norepinephrine depletion using DSP4 or global knockout of dopamine-beta hydroxylase (therefore reducing norepinephrine levels) have each been shown to reduce active avoidance learning (46–48,58,59). Additionally, the BNST specifically appears to play a similar role in biasing threat responses towards active coping behaviors, including active struggle bouts during restraint stress, which are associated with increased BNST neural activity (60). Further, chemogenetically inhibition of BNST neurons reduces active avoidance shuttling responses (61). While our data are consistent with the hypothesis that the vagal afferent-to-cNTS^A2^-to-vlBNST circuit is necessary for active responses to threat, additional work will be needed to examine the role of gut-sensing vagal afferents and noradrenergic inputs to the vlBNST in active avoidance behavior.

The data presented here are the first to examine retrieval of conditioned passive avoidance behavior specifically in rats in which gut-sensing (*Cckar*-expressing) vagal sensory afferents or noradrenergic inputs to the vlBNST were lesioned after training had already occurred. The temporal and circuit specificity of these lesions allowed us to isolate our analysis on the role of these circuit components in passive avoidance memory retrieval, without interfering with earlier phases of learning and memory. It is important to note, however, that while these lesions were made after footshock training, they permanently destroyed the affected neuronal populations. Thus, compensatory neuroadaptations could have occurred during the two week post-lesion period. In addition, the effects of partial chronic lesions such as those in the present study could differ from the effects of more transient or more complete interruption of vagal sensory and/or noradrenergic pathways. Future research should examine how more acute and reversible manipulations of neural activity in these ascending circuits influence passive avoidance retrieval.

In *Experiment 1, Cckar-*expressing neurons in the left nodose ganglia were more completely lesioned than *Cckar*+ cells in the right nodose ganglia. It is interesting to note that both preclinical and clinical studies as well as treatments utilizing VNS tend to administer stimulation unilaterally to the left vagus (62,63). While no studies have examined the effect of VNS on passive avoidance retrieval, left VNS has been shown to reduce figural memory retrieval (50) and to reduce symptoms of anxiety in some human patients (64,65). Since there is some indication of lateralized functions of the left vs. right vagus nerve in mice and rats (66,67), it would be interesting to examine whether laterality exists in vagal afferent pathways that modulate memory and avoidance.

Female rats were not included in our current study based on our previous findings that passive avoidance behavior is displayed more strongly in male rats, and that visceral feedback about metabolic state modulates passive avoidance in male but not female rats (3). Previous work has demonstrated sex-specific effects of central norepinephrine depletion on spatial memory retrieval in the Barnes maze, such that norepinephrine depletion improves performance in males, but impairs performance in females (54). Therefore, it will be important in future work to examine the role of vlBNST-projecting noradrenergic inputs on passive avoidance memory retrieval in females.

## Conclusions

Overall, results from the present study demonstrate that lesioning gut-sensing neurons in the nodose ganglia and noradrenergic neurons whose axons target the vlBNST enhances conditioned passive avoidance behavior in male rats. These new findings support the view that a gut vagal afferent-to-cNTS^NA^-to-vlBNST circuit plays a role in modulating the expression/retrieval of passive avoidance. Understanding how this circuitry contributes to conditioned avoidance may provide insight into effective treatment strategies for excessive avoidance behaviors in human anxiety and stress-related disorders.

## Declarations

### Consent for publication

Not applicable.

### Availability of data and materials

All data generated or analyzed during this study are included in this published article. Further enquiries can be directed to the corresponding author.

### Authors’ contributions

CME and LR designed the experiments. CME, IEG, DT, and TD performed experiments. CME and LR wrote the manuscript. All authors read and approved the final manuscript.

## Acknowledgements

Authors would like to thank Dr. Huiyuan Zheng for technical training and expertise and Dr. Diana Williams for use of equipment.

## List of Abbreviations

cNTS: caudal nucleus of the solitary tract
CSAP: cholecystokinin-conjugated saporin toxin DbH dopamine-beta hydroxylase
DSAP: dopamine-beta hydroxylase antibody-conjugated saporin toxin LC locus coeruleus
NA: noradrenergic
ROI: region of interest
vlBNST: ventrolateral bed nucleus of the stria terminalis VLM ventrolateral medulla
VNS: vagal nerve stimulation

